# Drug Target Ontology to Classify and Integrate Drug Discovery Data

**DOI:** 10.1101/117564

**Authors:** Yu Lin, Saurabh Mehta, Hande Küçük-McGinty, John Paul Turner, Dusica Vidovic, Michele Forlin, Amar Koleti, Dac-Trung Nguyen, Lars Juhl Jensen, Rajarshi Guha, Stephen L. Mathias, Oleg Ursu, Vasileios Stathias, Jianbin Duan, Nooshin Nabizadeh, Caty Chung, Christopher Mader, Ubbo Visser, Jeremy J. Yang, Cristian G. Bologa, Tudor Oprea, Stephan C. Schürer

## Abstract

**Background:** One of the most successful approaches to develop new small molecule therapeutics has been to start from a validated druggable protein target. However, only a small subset of potentially druggable targets has attracted significant research and development resources. The Illuminating the Druggable Genome (IDG) project develops resources to catalyze the development of likely targetable, yet currently understudied prospective drug targets. A central component of the IDG program is a comprehensive knowledge resource of the druggable genome.

**Results:** As part of that effort, we have been developing a framework to integrate, navigate, and analyze drug discovery data based on formalized and standardized classifications and annotations of druggable protein targets, the Drug Target Ontology (DTO). DTO was constructed by extensive curation and consolidation of various resources. DTO classifies the four major drug target protein families, GPCRs, kinases, ion channels and nuclear receptors, based on phylogenecity, function, target development level, disease association, tissue expression, chemical ligand and substrate characteristics, and target-family specific characteristics. The formal ontology was built using a new software tool to auto-generate most axioms from a database while also supporting manual knowledge acquisition. A modular, hierarchical implementation facilitates development and maintenance and makes use of various external ontologies, thus integrating the DTO into the ecosystem of biomedical ontologies. As a formal OWL-DL ontology, DTO contains asserted and inferred axioms. Modeling data from the Library of Integrated Network-based Cellular Signatures (LINCS) program illustrates the potential of DTO for contextual data integration and nuanced definition of important drug target characteristics. DTO has been implemented in the IDG user interface Portal, Pharos and the TIN-X explorer of protein target disease relationships.

**Conclusions:** DTO was built based on the need for a formal semantic model for druggable targets including various related information such as protein, gene, protein domain, protein structure, binding site, small molecule drug, mechanism of action, protein tissue localization, disease association, and many other types of information. DTO will further facilitate the otherwise challenging integration and formal linking to biological assays, phenotypes, disease models, drug poly-pharmacology, binding kinetics and many other processes, functions and qualities that are at the core of drug discovery. The first version of DTO is publically available via the website http://drugtargetontology.org/, Github (https://github.com/DrugTargetOntology/DTO), and the NCBO Bioportal (https://bioportal.bioontology.org/ontologies/DTO). The long-term goal of DTO is to provide such an integrative framework and to populate the ontology with this information as a community resource.

## 1 INTRODUCTION

The development and approval of novel small molecule therapeutics (drugs) is highly complex and exceedingly resource intensive, being estimated at over one billion dollars for a new FDA approved drug. The primary reason for attrition in clinical trials is the lack of efficacy, which has been associated with poor or biased target selection [1]. Although the drug target mechanism of action is not required for FDA approval, a target-based mechanistic understanding of diseases and drug action is highly desirable and a preferred approach of drug development in the pharmaceutical industry. Following the advent of the Human Genome, several research groups in academia as well as industry have focused on “the druggable genome” *i.e*. the subsets of genes in the human genome that express proteins which have the ability to bind drug-like small molecules [2]. The researchers have estimated the number of druggable targets ranging from few hundred to several thousands [3]. Furthermore, it has been suggested by several analyses that only a small fraction of likely relevant druggable targets are extensively studied, leaving a potentially huge treasure trove of promising, yet understudied (“dark”) drug targets to be explored by pharmaceutical companies and academic drug discovery researchers. Not only is there ambiguity about the number of the druggable targets, but there is also a need of systematic characterization and annotation of the druggable genome. A few research groups have made efforts to address these issues and have indeed developed several useful resources, *e.g*. IUPHAR/BPS Guide to PHARMACOLOGY (GtoPdb/IUPHAR) [4], PANTHER [5], Therapeutic Target Database (TTD) [6], Potential Drug Target Database (PDTD) [7], covering important aspects of the drug targets. However, to the best of our knowledge, a publically available structured knowledge resource of drug target classifications and relevant annotations for the most important protein families, one that facilitates querying, data integration, re-use, and analysis does not currently exist. Content in the above-mentioned databases is scattered and in some cases inconsistent and duplicated, complicating data integration and analysis.

The Illuminating the Druggable Genome (IDG) project (http://targetcentral.ws/) has the goal to identify and prioritize new prospective drug targets among likely targetable, yet currently poorly or not at all annotated proteins; and by doing so to catalyze the development of novel drugs with new mechanism of action. Data compiled and analyzed by the IDG Knowledge Management Center (IDG-KMC) shows that the globally marketed drugs stem from only 3% of the human proteome. These results also suggest that the substantial knowledge deficit for understudied drug targets may be due to an uneven distribution of information and resources [8].

In the context of the IDG program we have been developing the Drug Target Ontology (DTO). Formal ontologies have been quite useful to facilitate harmonization, integration, and analysis of diverse data in the biomedical and other domains. DTO integrates and harmonizes knowledge of the most important druggable protein families: kinases, GPCRs, ion channels and nuclear hormone receptors. DTO content was curated from many information sources and the literature and includes detailed hierarchical classifications of proteins and genes, tissue localization, disease association, drug target development level, protein domain information, ligands, substrates, and other types of relevant information. The covered resources were chosen by domain experts based on relevance coverage and completeness of the information available through them. Most resources had been peer reviewed (references are included at appropriate places), published and were therefore considered reliable. DTO is aimed towards the drug discovery and clinical communities and was built to align with other ontologies including BioAssay Ontology (BAO) [9-11] and GPCR Ontology [12]. By providing a semantic framework of diverse information related to druggable proteins, DTO facilitates the otherwise challenging integration and formal linking of heterogeneous and diverse data important for drug discovery. DTO is particularly relevant for big data, systems-level models of diseases and drug action as well as precision medicine. The long-term goal of DTO is to provide such an integrative framework and to populate the ontology with this information as a community resource. Here we describe the development, content, architecture, modeling and use of the DTO. DTO has already been implemented in end-user software tools to facilitate the browsing [11] and navigation of drug target data [13].

## 2 Methods

### 2.1 Drug Target data curation and classification

DTO places special emphasis on the four protein families that are central to the NIH IDG initiative: non-olfactory GPCRs (oGPCRs), Kinases, Ion Channels and Nuclear Receptors. The classifications and annotations of these four protein families were extracted, aggregated, harmonized, and manually curated from various resources as described below, and further enriched using the recent research literature. Proteins and their classification and annotations were aligned with the Target Central Resource Databases (TCRD) database [11] developed by the IDG project (http://targetcentral.ws/ProteinFam). In particular, the Target Development Level (TDL) classification was obtained from the TCRD database.

#### Kinase classification

Kinases have been classified primarily into protein and non-protein kinases. Protein kinases have been further classified into several groups, families, subfamilies. Non-protein kinases have been classified in several groups, based on the type of substrates (lipid, carbohydrate, nucleoside, other small molecule, *etc*.). Classification information has been extracted and curated from various resources *e.g*. UniProt, ChEMBL, PhosphoSitePlus^®^ (PSP) [14], Sugen Kinase website (http://www.kinase.com/web/current/), and the literature, and was organized manually, consolidated and checked for consistency. Kinase substrates were manually curated from UniProt and the literature. Pseudokinases, which lack key functional residues and are (to current knowledge) not catalytically active, were annotated based on the Sugen kinase domain sequences and the literature.

#### Ion-channel classification

Ion channels have been classified primarily into family, subfamily, sub-subfamily. Most of the information has been taken from the Transporter Classification Database (http://www.tcdb.org/) [15], UniProt and several linked databases therein. The classification is based on both the phylogenetic and functional information. Additional information regarding the gating mechanism (voltage gated, ligand gated, etc.), transported ions, protein structural and topological information has also been captured and included as separate annotations. Moreover, the transported ions, such as chloride, sodium, etc. have been mapped to the “Chemical entity” of the ChEBI reference database [16].

#### GPCR classification

GPCRs have been classified based on phylogenetic, functional and the endogenous ligand information. The primary classification included class, group, family, and subfamily. Most of the information has been taken from the GPCR.org classification and had been updated using various sources *e.g*. IUPHAR [4], ChEMBL, UniProt and also from our earlier GPCR ontology [12]. Furthermore, the information for the specific endogenous ligands for each protein has been extracted from IUPHAR and has been integrated with the classification. The information about the GPCR ligand and ligand type (lipid, peptide, *etc*.) has also been included and has been mapped manually to the “Chemical entity” of the ChEBI reference database.

#### Nuclear receptor classification

This information has been adopted directly from IUPHAR.

#### External DTO modules and mapping: Proteins mapped to UniProt

Genes were classified identical to proteins (above) and mapped to Entrez gene. The external modules incorporated into DTO were extracted from the Disease Ontology (DOID) [17], BRENDA Tissue Ontology (BTO) [18], UBERON [19], the ontology of Chemical Entities of Biological Interest (ChEBI) [20], and Protein Ontology (PRO) [21]. Data about over 1000 cell lines from the LINCS project [22] were integrated and mapped to diseases and tissues. Gene/protein–disease [23] and protein–tissue associations [24] were obtained from the JensenLab at Novo Nordisk Foundation Center for Protein Research. Mapping between UBERON and BRENDA to integrate the tissue associations of cell lines and proteins was retrieved from the NCBO BioPortal [25, 26] and manually cross-checked. Target Development Level (TDL) were obtained from TCRD and included as separate annotation for all protein families.

### 2.2 Drug Target Ontology (DTO) development

#### Ontology modeling

While curators stored all classification and annotation data into various spreadsheets, ontologists created the ontological model to link the metadata obtained from those spreadsheets, and to create the descriptive logic axioms to define ontology classes using a semi automated workflow. Finalizing and optimizing the ontology model or design pattern required iterative processes of intensive discussions, modeling refinement, voting, and approval among domain experts, data curators, IT developers, and ontologists. Once ontologists proposed a conceptual ontology model, the selection of the most robust ontology model was guided by simple criteria: correct representation of domain content, minimize the number of relations to link all metadata, avoid contradiction with existing domain knowledge representation ontologies, such as the OBO ontologies. For example, in our conceptual model, the relations among organ, tissue, cell lines and anatomical entity were adopted and refined from the UBERON and CLO ontologies. Some relations such as the shortcut relations between protein and associated disease or tissue were created specifically for DTO, which was a compromise for accommodating the large amount of data in DTO. Approval process of accepting a model proposal was driven by our domain experts with contributing data curators, IT developers, and ontologists. The voting process was rather informal; however, the model had to be agreed by all the parties involved in the ontology development: domain experts, data curators, IT developers, and ontologists. Once the most fit ontology model was chosen, this piece of modeling was used as template for a java tool (described below) to generate all the OWL files by using above mentioned data annotation spreadsheets as input.

#### Modularization approach

DTO was built with an extended modular architecture based on the modular architecture designed and implemented for BAO [9]. The modularization strategy developed previously was a layered architecture and used the modeling primitives, vocabularies, modules and axioms. Most significantly, DTO’s modular architecture includes an additional layer to the modularization process by automating the creation of basic subsumption hierarchies and select axioms such as the axioms for disease and tissue associations. Three types of files are used in the modular architecture: vocabulary files, module files, and combined files, such as DTO_core and DTO_complete. Vocabularies only contain concepts (classes with subsumption only). Module layers enable combining vocabularies in flexible ways to create desired ontology structures or subsets. Finally, in the combined files axioms are added to the vocabularies to formally define the various concepts to allow logical inferences. Classes and relationships are imported (directly or indirectly) from module and/or vocabulary files [9]. The external third party ontologies were extracted using the OWL API or OntoFox [27].

#### OntoJOG tool

To streamline the building process, a Java tool (OntoJOG) was developed to automatically create the OWL module files, vocabulary files as components of the whole ontology. OntoJOG takes a flat CSV or TSV data file and loads it as a table either into a temporary SQLite database or a permanent MySQL database. This table is then used as a reference for creating and generating the OWL files as well as several relationship tables. The relationship tables and the final OWL files are generated based on a CSV mapping file that generates the commands for the OntoJOG to perform and the various options for those commands. The commands from the mapping file are read in two passes to ensure everything is added correctly. In the first pass all classes and their annotations are inserted into the relationship tables and are assigned IDs as necessary, and in the second pass all axioms and relationships between classes are created. After this process is completed an optional reparenting phase is executed before each module of the ontology is generated into its own OWL vocabulary files with an accompanying module file containing the relationships for the given vocabulary files. Finally, the ontology was thoroughly reviewed, tested and validated by developers, domain experts, and users in the IDG-KMC.

### 2.3 DTO Visualization

Data visualization is important, especially with the increasing complexity of the data. Ontology visualization, correspondingly, has an appealing potential to help to browse and comprehend the structures of ontologies. A number of ontology visualization tools have been developed and applied as information retrieval aids, such as OntoGraf, OWLViz as part of the Ontology development tool Protégé, and OntoSphere3D [28] among others. Further, studies and reviews on different visualization tools, e.g. [29, 30] and [31], have been published by comparing each tool’s performances. Preference of visualization models depends on the type and query context of the visualized network, also on users’ needs.

Data-Drive Document (D3) is a relatively novel representation-transparent and dynamic approach to visualize data on the web. It is a modern interactive visualization tool available as a JavaScript library [29]. By selectively binding input data to arbitrary document elements, D3.js enables direct inspection and manipulation of a native representation. The D3.js JavaScript library gained popularity as a generic framework based on widely accepted web standards such as SVG, JavaScript, HTML5 and CSS.

Consequently, we use the D3.js library for the interactive visualization of our DTO as part of the Neo4J graphical database solution.

### 2.4 DTO and BAO integration to model LINCS data

The Library of Network-Based Cellular Signatures (LINCS) program has been generating a reference “library” of molecular signatures, such as changes in gene expression and other cellular phenotypes that occur when cells are exposed to a variety of perturbing agents. One of the LINCS screening assays is a biochemical kinase profiling assay that measures drug binding using a panel of ~440 recombinant purified kinases, namely, KINOMEscan assay. The HMS LINCS Center has collected 165 KINOMEscan datasets in order to analyze the drug-target interaction. All these LINCS KINOMEscan data were originally retrieved from Harvard Medical School (HMS) LINCS DB (http://lincs.hms.harvard.edu/db/). KINOMEscan data was curated by domain experts to map to both Pfam domains, and corresponding Kinases. Unique KINOMEscan domains and annotations, including domain descriptions, IDs, names, gene symbols, phosphorylation status, and mutations were curated from different sources, including the HMS LINCS DB, DiscoverX KINOMEscan^®^ assay list [32], Pfam (http://pfam.xfam.org/), and our previous modeling efforts of the entire human Kinome (publication in preparation). The kinase domain classification into group, family, *etc*. was the same as described above (kinase classification). Gatekeeper and hinge residues were assigned based on structural alignment of existing kinase domain crystal structures and structural models of the human kinome and sequence alignment with the full kinase protein referenced by UniProt accession in the DTO. Pfam accession number and names were obtained from Pfam [33]. The protocol and the KINOMEscan curated target metadata table were analyzed by ontologists to create kinase domain drug target ontology model.

### 2.5 Ontology source access and license

The official DTO website is publicly available at http://drugtargetontology.org/, where it can be visualized and searched. The DTO is an open source project, and released under a Creative Commons 3.0 License. The source code including the development and release versions are freely available at the URL: http://github.com/DrugTargetOntology/DTO. DTO is also published at the NCBO BioPortal (https://bioportal.bioontology.org/ontologies/DTO).

## 3 Results

In what follows, the *italic* font represents terms, classes, relations, or axioms used in the ontology.

### 3.1 Drug Targets definition and classification

Different communities have been using the term “drug target” ambiguously with no formal generally accepted definition. The DTO project develops a formal semantic model for drug targets including various related information such as protein, gene, protein domain, protein structure, binding site, small molecule drug, mechanism of action, protein tissue localization, disease associations, and many other types of information.

The IDG project defines ‘drug target’ as “a native (gene product) protein or protein complex that physically interacts with a therapeutic drug (with some binding affinity) and where this physical interaction is (at least partially) the cause of a (detectable) clinical effect”. DTO defined a DTO specific term “*drug target role*”. The text definition of “*drug target role*” is “a role played by a material entity, such as native (gene product) protein, protein complex, microorganism, DNA, etc., that physically interacts with a therapeutic or prophylactic drug (with some binding affinity) and where this physical interaction is (at least partially) the cause of a (detectable) clinical effect.” At the current phase, DTO focuses on protein targets. DTO provides various asserted and inferred hierarchies to classify drug targets. Below we describe the most relevant ones.

#### 3.1.1 Target Development Level (TDL)

The IDG classified proteins into four levels with respect to the depth of investigation from a clinical, biological and chemical standpoint (http://targetcentral.ws/) [8]:

1. **T_clin_** are proteins targeted by approved drugs as they exert their mode of action [3]. The Tclin proteins are designated drug targets under the context of IDG.
2. **T_chem_** are proteins that can specifically be manipulated with small molecules better than bioactivity cutoff values (30 nM for kinases, 100 nM for GPCRs and NRs, 10 uM for ICs, and 1 uM for other target classes), which lack approved small molecule or biologic drugs. In some cases, targets have been manually migrated to Tchem through human curation, based on small molecule activities from sources other than ChEMBL or Drug Central.
3. **T_bio_** are proteins that do not satisfy the T_clin_ or T_chem_ criteria, which are annotated with a Gene Ontology Molecular Function or Biological Process with an Experimental Evidence code, or targets with confirmed OMIM phenotype(s), or do not satisfy the Tdark criteria detailed in 4).
4. **T_dark_** refers to proteins that have been described at the sequence level and have very few associated studies. They do not have any known drug or small molecule activities that satisfy the activity thresholds detailed in 2), lack OMIM and GO terms that would match Tbio criteria, and meet at least two of the following conditions:

- A PubMed text-mining score < 5 [23]
- <= 3 Gene RIFs [34]
- <= 50 Antibodies available per Antibodypedia (http://antibodypedia.com)

#### 3.1.2 Functional and phylogenetic classification

DTO proteins have been classified into various categories based on their structural (sequence/domains) or functional similarity. A high-level summary of the classifications for Kinases Ion Channels, GPCRs and Nuclear Receptors is shown in Figure 1. It should be noted that, as indicated above, the classification information has been extracted from various database and literature resources. The classification is subject to continuous updating for greater accuracy, and enriching the DTO using the most recent information as it becomes available. The present classification of the four protein families is briefly discussed below:

**Figure 1.**
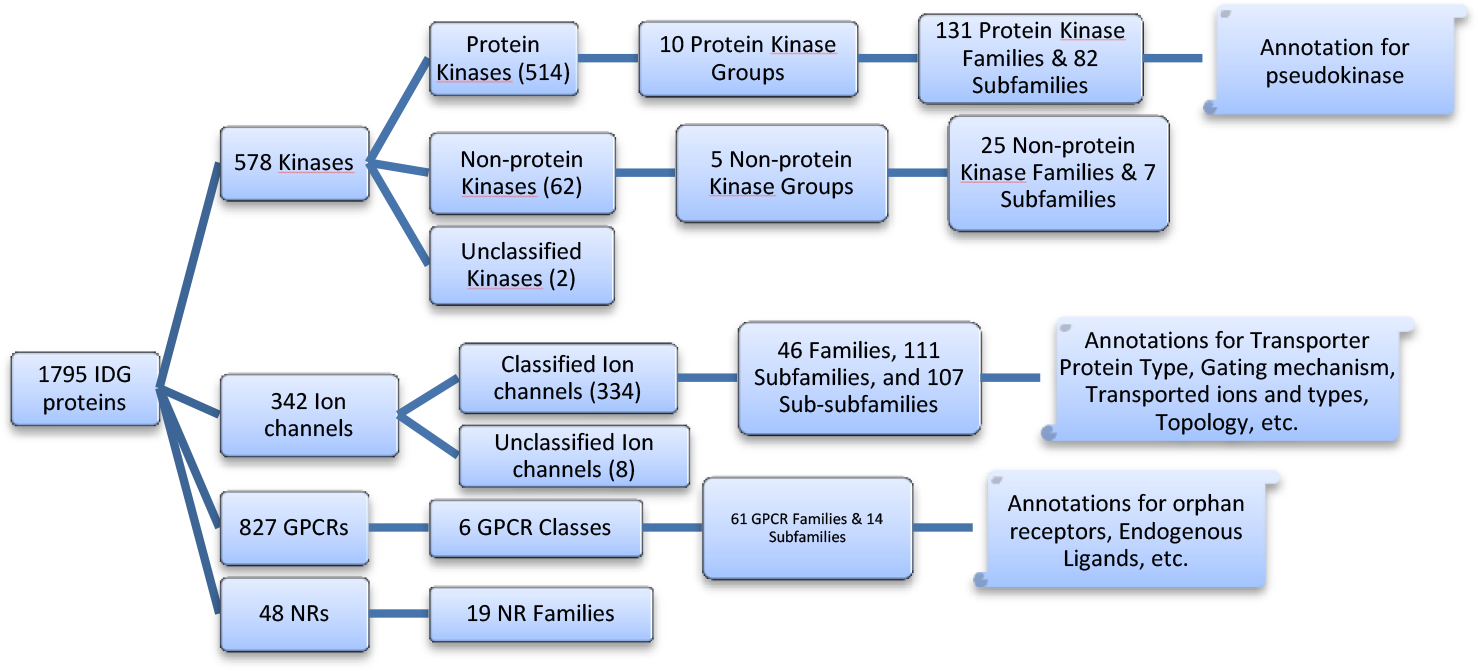
Summary of protein classification hierarchies for Kinase, Ion Channel, GPCR, and NR protein families. Note that several other relevant target annotations have been incorporated into DTO.

Most of the 578 kinases covered in the current version of DTO are protein kinases. These 514 PKs are categorized in 10 groups that are further sub-categorized in 131 families and 82 subfamilies. A representative classification hierarchy for MAPK1 is:

*Kinase > Protein Kinase > CMGC group > MAPK family > ERK subfamily > Mitogen-activated Protein Kinase 1*

The 62 non-protein kinases are categorized in 5 groups depending upon the substrate that are phosphorylated by these proteins. These 5 groups are further sub-categorized in 25 families and 7 subfamilies. There are two kinases that haven’t been categorized yet in any of the above types or groups.

The 334 Ion channel proteins (out of 342 covered in the current version of DTO) are categorized in 46 families, 111 subfamilies, and 107 sub-subfamilies.

Similarly, the 827 GPCRs covered in the current version of DTO are categorized in 6 classes, 61 families and 14 subfamilies. The additional information whether any receptor has a known endogenous ligand or is currently “orphan” is mapped with the individual proteins. Finally, the 48 nuclear hormone receptors are categorized in 19 NR families.

#### 3.1.3 Disease- and tissue-based classification

Target-disease associations and tissue expressions were obtained from the DISEASES [23]and TISSUES [24] databases (see Methods). Examples of such classifications are available as inferences in DTO (see below section 3.3.2).

#### 3.1.4 Additional annotations and classifications

In addition to the phylogenetic classification of the proteins, there are several relevant properties associated with them as additional annotations. For example, there are 46 PKs that have been annotated as pseudokinases [35]. For ion channels, important properties, like transporter protein type, transported ion(s), gating mechanism, etc. have been associated with the individual proteins. The gating mechanism refers to the information regarding the factors that control the opening and closing of the ion channels. The important mechanisms include voltage-gated, ligand-gated, temperature-gated, mechanically-gated, etc. Similarly, for the GPCRs, the additional information whether any receptor has a known endogenous ligand or is currently “orphan” is mapped with the individual proteins. Current version of DTO has approximately 255 receptors that have information available regarding the endogenous ligands.

The analysis of Drug target protein classification along with such relevant information associated through separate annotations may lead to interesting inferences.

#### 3.1.5 Chemical classifications

Known GPCR ligands and IC transported ions were categorized by chemical properties and mapped to ChEBI (see methods). For example, depending upon their chemical structure and properties, these known endogenous ligands for GPCRs have been categorized in seven types, namely, amine, amino acid, carboxylic acid, lipid, peptide, nucleoside and nucleotide. Similarly, the ions transported by the ion channel proteins and ion types (anion/cation) have been mapped to ChEBI. These annotations together with mappings of substrates and ligands to the proteins enable inferred classification of the proteins based on their chemical properties (see below).

### 3.2 DTO ontology implementation and modeling

#### 3.2.1 Drug discovery target knowledge model of the DTO

The first version of the DTO includes detailed target classification and annotations for the four IDG protein families. Each protein is related to four types of entities: gene, related disease, related tissue or organ, and target development level. The conceptual model of DTO is illustrated as a linked diagram with nodes and edges. Nodes represent the classes in the DTO, and edges represent the ontological relations between classes. As shown in Figure 2, GPCRs, kinases, ICs and NRs are types of proteins. GPCR binds GPCR ligands, and IC transports ions. Most GPCR ligands and ion are types of chemical entity from ChEBI. Each protein has a target development level (TDL), i.e., T_clin_, T_chem_, T_bio_ and T_dark_. The protein is linked to gene by ‘has gene template’ relation. The gene is associated with disease based on evidence from the DISEASES database. The protein is also associated with some organ, tissue, or cell line using some evidence from TISSUES database. The full DTO contains many more annotations and classifications, and it is available at http://drugtargetontology.org/

**Figure 2.**
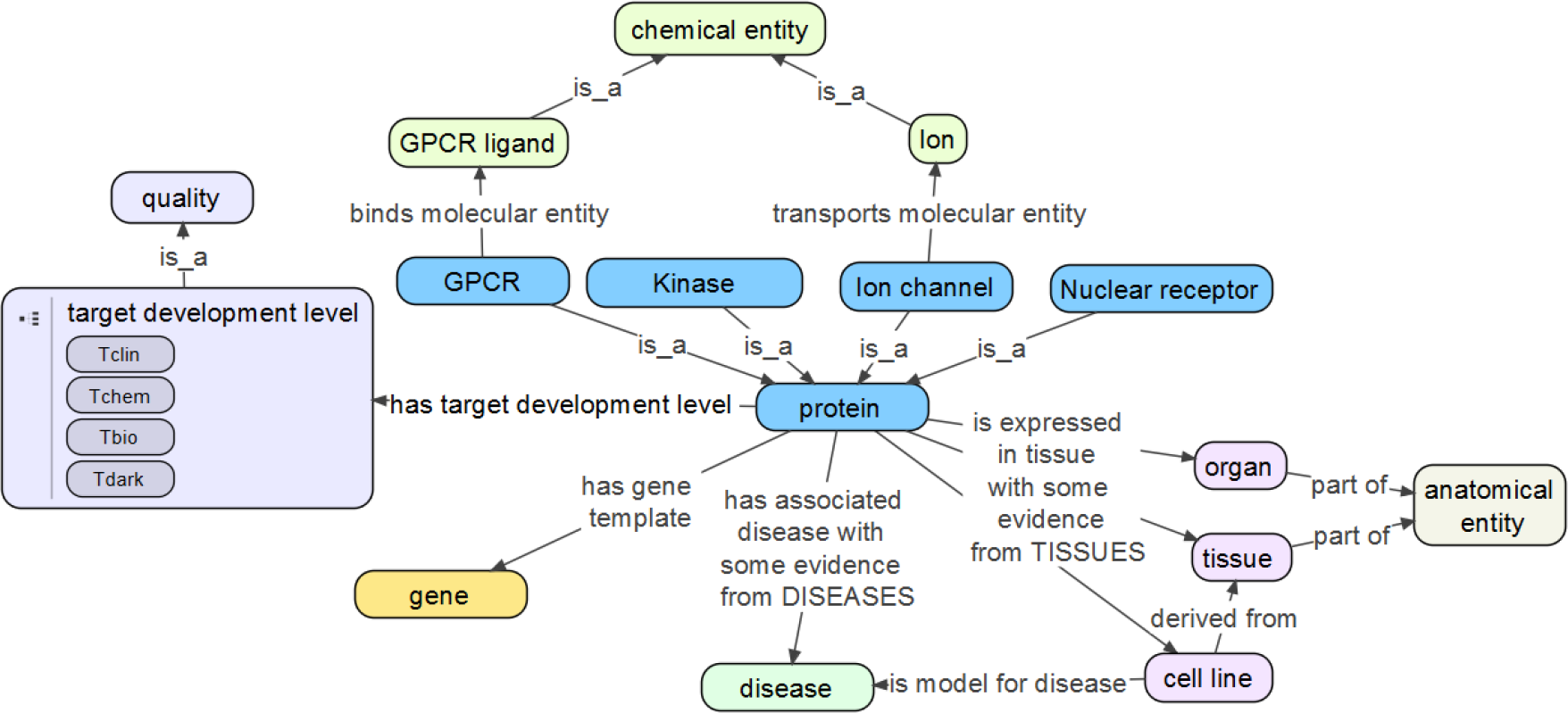
Conceptual high-level model of main DTO entities.

DTO is implemented in OWL2-DL to enable further classification by inference reasoning and SPARQL queries. The current version of DTO contains > 13,000 classes and > 220,000 axioms. The DTO contains 827 GPCRs, 572 kinase, 342 ion channels (ICs), and 48 NRs.

#### 3.2.2 Modular implementation of the DTO combining auto-generated and expert axioms

In DTO, each of the four drug target families has two vocabulary files of gene and protein, respectively; other DTO-native categories were created as separate vocabulary files. Additional vocabulary files include quality, role, properties, and cell line classes and subclasses. A vocabulary file contains entities of a class, which only contains “is-a” hierarchies. For example, the GPCR gene vocabulary contains only GPCR gene list and its curated classification. DTO core imports all the DTO vocabulary files of four families, including genes and proteins, and necessary axioms were added. Finally, DTO core was imported into the DTO complete file, which includes other vocabulary files and external files. External ontologies used in DTO include: BTO, CHEBI, DOID, UBERON, Cell Line Ontology (CLO), Protein Ontology (PRO), Relations Ontology (RO) and Basic Formal Ontology (BFO). The DTO core and DTO external are imported into the DTO module with auto-generated axioms, which links entities from different vocabulary files. Besides the programmatically generated vocabularies and modules, DTO also contains manually generated vocabularies and modules, as shown in Figure 3.

**Figure 3.**
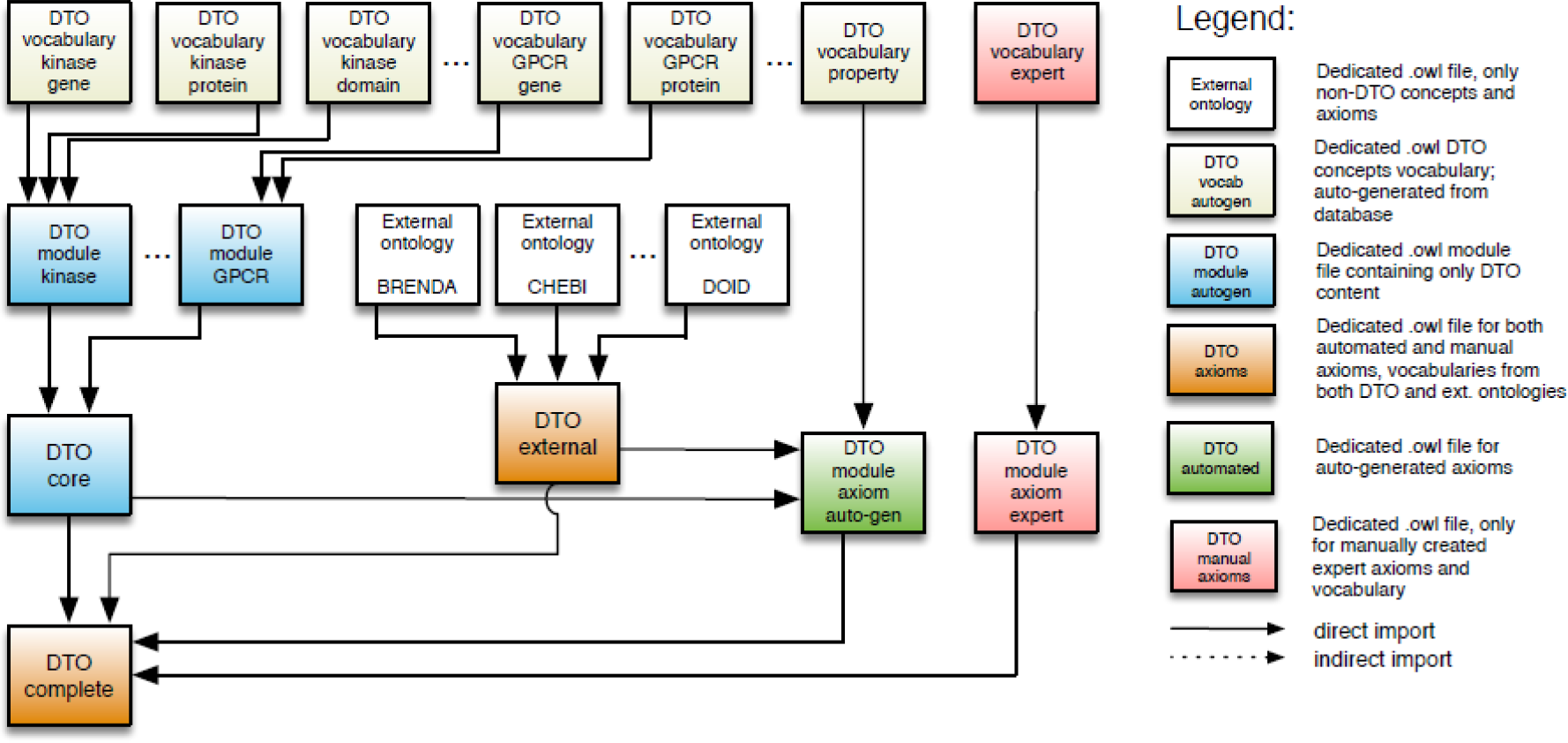
DTO architecture.

This approach significantly simplifies the maintenance of the ontology contents, especially when the ontology is large in size. If the gene or protein list changes, only the vocabulary file and the specific module file need to be updated instead of the whole ontology. In addition, external and internal resources are maintained separately. This modularization approach facilitates automated content updates from external resources including axioms generated using the above-mentioned Java tool OntoJOG without the need to re-generate manually axiomized domain knowledge, which can be very resource intensive, by simply separating them into two layers.

### 3.3 DTO to infer biologically and chemically relevant target classes

#### 3.3.1 Chemically relevant target classes inferred by DTO

In addition to detailed asserted target classifications, DTO incorporates various other annotations including GPCR endogenous ligands for GPCRs, transported ions for ICs, gating mechanism for ICs, or pseudokinases. Endogenous GPCR ligands were manually mapped to ChEBI and classified by chemical category such as amine, lipid, peptide, etc. As ligands relate to receptor properties, GPCRs are typically classified based on their ligands; however, the ligand-based classification is orthogonal to the classification based on class A, B, C, adhesion, etc. and it changes as new ligands are deorphanized.

In DTO we therefore infer the ligand-based receptor, for example *aminergic GPCR*, *lipidergic GPCR*, *peptidic GPCR*, and *orphan GPCR*, which are of particular interest, by defining their logical equivalent as follows:

*aminergic GPCR ≡ GPCR and (’binds molecular entity’ some amine);*

*lipidergic GPCR ≡ GPCR and (’binds molecular entity’ some lipid);*

*peptidic GPCR ≡ GPCR and (’binds molecular entity’ some peptide);*

*orphan GPCR ≡ GPCR and (not (’binds molecular entity’ some ‘GPCR ligand’))*.

An example for 5-hydroxytryptamine receptor is shown in Figure 4; the receptor is inferred as aminergic receptor based on its endogenous ligand.

**Figure 4.**
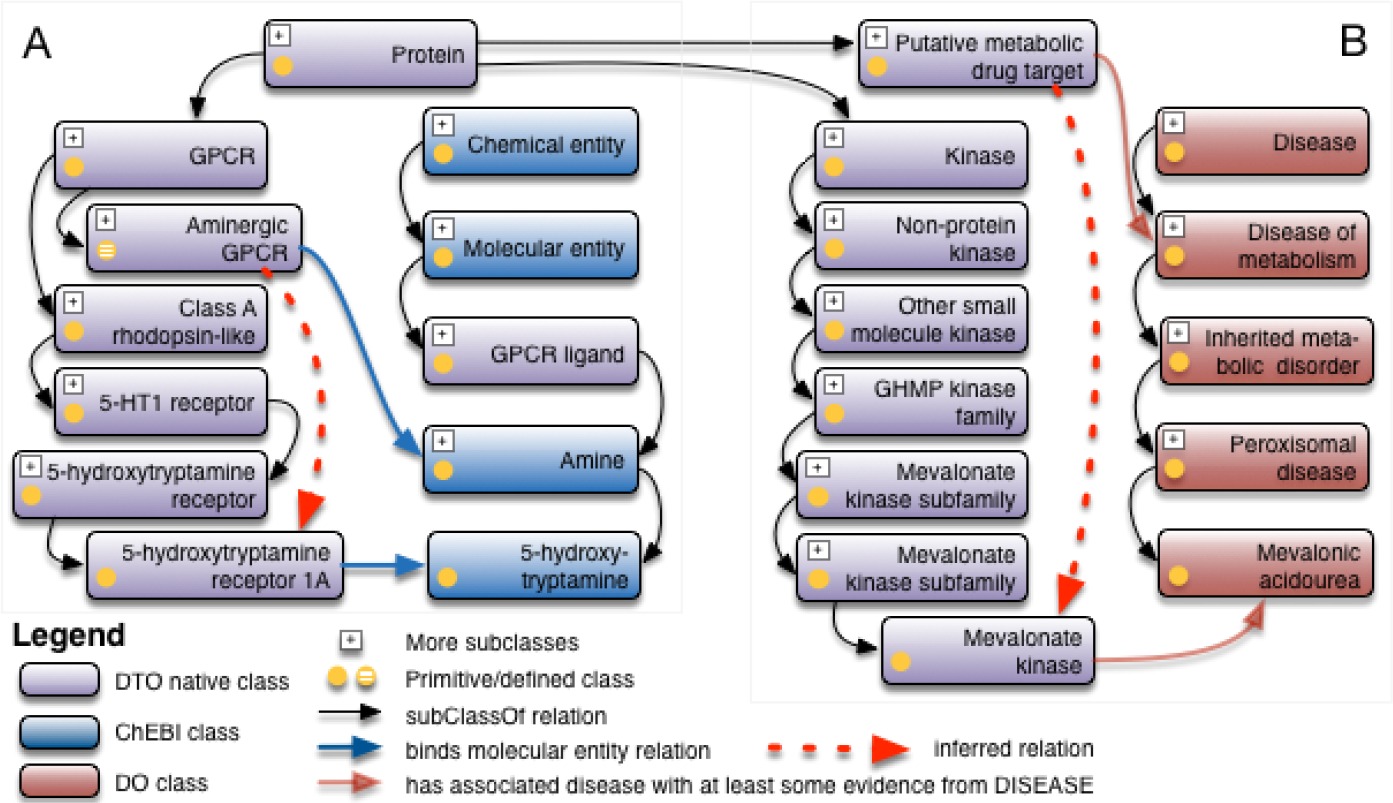
Inferred classifications in DTO: 5-hydroxytryptamine receptor as aminergic receptor based in its endogenous ligand, mevalonate kinase as putative metabolic drug target.

DTO has classified 39 *aminergic GPCR*, 37 *lipidergic GPCR*, 119 *peptide GPCR* and 582 *orphan GPCR*.

#### 3.3.2 Disease relevant target classes inferred by DTO

In a similar way, we categorized important disease targets by inference based on the protein disease association, which were modeled as ‘strong’, ‘at least some’, or ‘at least weak’ evidence using subsumption. For example, DTO uses the following hierarchical relations to declare the relation between a protein and the associated disease extracted from the DISEASES database. *has associated disease with at least weak evidence from DISEASES*

*has associated disease with at least some evidence from DISEASES*
*has associated disease with strong evidence from DISEASES*

In the DISEASES database, the associated disease and protein are measured by a Z-Score [23]. In DTO, the “at least weak evidence” is translated as a Z-Score between zero and 2.4; the “some evidence” is translated as a Z-Score between 2.5 and 3.5; and the “strong evidence” is translated as a Z-Score between 3.6 and 5.

This allows querying or inferring proteins for a disease of interest by evidence. Diseases related targets were defined using following axioms (as illustrative as examples):

*Putative infectious disease targets ≡ Protein and (’has associated disease with strong evidence from DISEASES’ some ‘disease of metabolism’);*

*Putative infectious disease targets ≡ Protein and (’has associated disease with strong evidence from DISEASES’ some ‘disease by infectious agent’);*

*Putative mental health disease targets≡ Protein and (’has associated disease with strong evidence from DISEASES’ some ‘developmental disorder of mental health’)*

We created such inference examples in DTO, including 29 *metabolic disease targets*, 36 *mental health disease targets*, and 1 *infectious disease targets*.

### 3.4 Modeling and integration of Kinase data from the LINCS project

The Library of Network-Based Cellular Signatures (LINCS, http://lincsproject.org/) program has a systems biology focus. This project has been generating a reference “library” of molecular signatures, such as changes in gene expression and other cellular phenotypes that occur when cells are exposed to a variety of perturbing agents. The project also builds computational tools for data integration, access, and analysis. Dimensions of LINCS signatures include the biological model system (cell type), the perturbation (e.g. small molecules) and the assays that generate diverse phenotypic profiles. LINCS aims to create a full data matrix by coordinating cell types and perturbations as well as informatics and analytics tools. We have processed various LINCS datasets, which are available at the LINCS Data Portal (http://lincsportal.ccs.miami.edu/).

LINCS data standards [22]are the foundation of LINCS data integration and analysis. We have previously illustrated how integrated LINCS data can be used to characterize drug action [36]; among those, KINOME-wide drug profiling datasets.

We have annotated the KINOMEscan domains data generated from HMS LINCS KINOMEscan dataset. The annotation includes domains descriptions, names, gene symbols, phosphorylation status, and mutations. To integrate this information into DTO, we built a kinase domain module following the modularization approach described in section 2.2.

We started with an example scenario given by domain expert shown below:

ABL1 is a tyrosine-protein kinase with UNIPROT ID P00519 (human). The sequence itself is 1131 AA long.
The KINOMEscan domain named "ABL1" is a part of the protein (AA Start/Stop S229/K512) containing the "Pkinase-Tyr" domain (pFam accession PF07714.14, AA Start/Stop I242/F493)
The KINOMEscan domain named "ABL1(F317I)-nonphosphorylated" is the same part of the protein (AA Start/Stop S229/K512) with a mutation at position 317. pFam, anyway identifies the same domain (accession PF07714.14)
The KINOMEscan domain named "ABL1(F317I)-phosphorylated" is the same part of the protein (AA Start/Stop S229/K512) with a mutation at position 317. pFam, anyway identifies the same domain (accession PF07714.14)

In this scenario, there are four major ontological considerations or relations need to be considered when building an ontology module (Figure 5).

**Figure 5.**
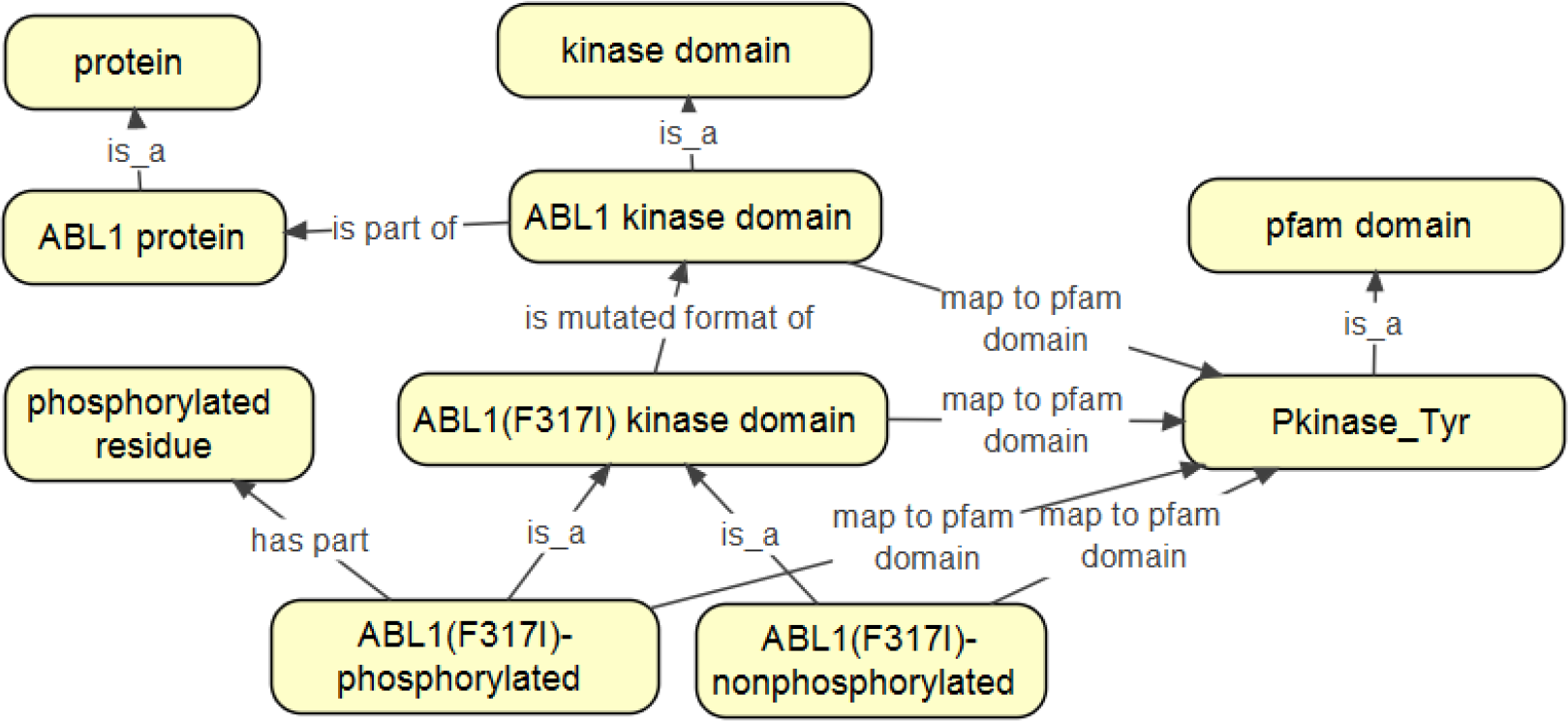
Relations between protein, kinase domain, mutated kinase domain, phosphorylated kinase domain, and pfam domains in DTO.

#### 3.4.1 Kinase domain and kinase protein

DTO use “*has part*” relation to link the kinase protein and kinase domain, which reflects the biological reality that kinase domain is a partial protein.

#### 3.4.2 Kinase domain variations: mutated kinase domain and phosphorylated kinase domain

Mutated kinase domain relates to its wild type kinase domain by simply using “*is mutated form of*” relation. Both, phosphorylated and nonphosphorylated forms of a kinase domain are child of a kinase domain from which they were modified to current phosphorylation variation forms. Since the KINOMEscan assay cannot provide the specific phosphorylation position information, the definition of a phosphorylated form of a kinase domain, either mutated or wild-type, is generally constituted using an ad-hoc axiom: *has part some “phosphorylated residue”*. Note that “*phosphorylated residue*” (MOD_00696) is an external class imported from Protein Modification Ontology (MOD).

#### 3.4.3 Pfam domain mapping to kinase domain and its variations

DTO data curators / domain experts have mapped all the kinase domains (including their variations) to Pfam families using sequence level data. This information was captured by using “*map to pfam domain*” relation, which links a kinase domain to a pfam domain.

Figure 5 shows how in DTO the above scenario is modeled by connecting *ABL1 Kinase domain* with *ABL1 protein* using relation *is part of*, as well as how kinase domain relates to Pfam domain using *map to pfam domain* relation. In this scenario, all the variations of ABL1 kinase domain are mapped to the same Pfam domain.

#### 3.4.4 Kinase gatekeeper and mutated amino acid residues

The kinase gatekeeper position is an important recognition and selectivity element for small molecule binding. One of the mechanisms by which cancers evade kinase drug therapy is by mutation of key amino acids in the kinase domain. Often the gatekeeper is mutated. Located in the ATP binding pocket of protein kinases, the gatekeeper residue has been shown to influence selectivity and sensitivity to a wide range of small molecule inhibitors. Kinases that possess a small side chain at this position (Thr, Ala, or Gly) are readily targeted by structurally diverse classes of inhibitors, whereas kinases that possess a larger residue at this position are broadly resistant [37].

DTO defines a “*gatekeeper role*” to capture any residue that is annotated as a gatekeeper. In the case of ABL1 kinase domain, the THR74 within the ABL1 kinase domain is identified as a gatekeeper by the data curator / domain expert. This gatekeeper residue is further mapped to the 315^th^ residue located in the whole ABL1 kinase amino acid sequence. DTO defines a term: *THR315 in ABL1 kinase domain* with an axiom of “*has role some gatekeeper role*”. With an equivalence definition of term “*gatekeeper residue*” as anything that satisfied the condition of “*has role some gatekeeper role*”, DTO can group all the gatekeeper residues in this KINOMEscan dataset (Figure 6).

**Figure 6.**
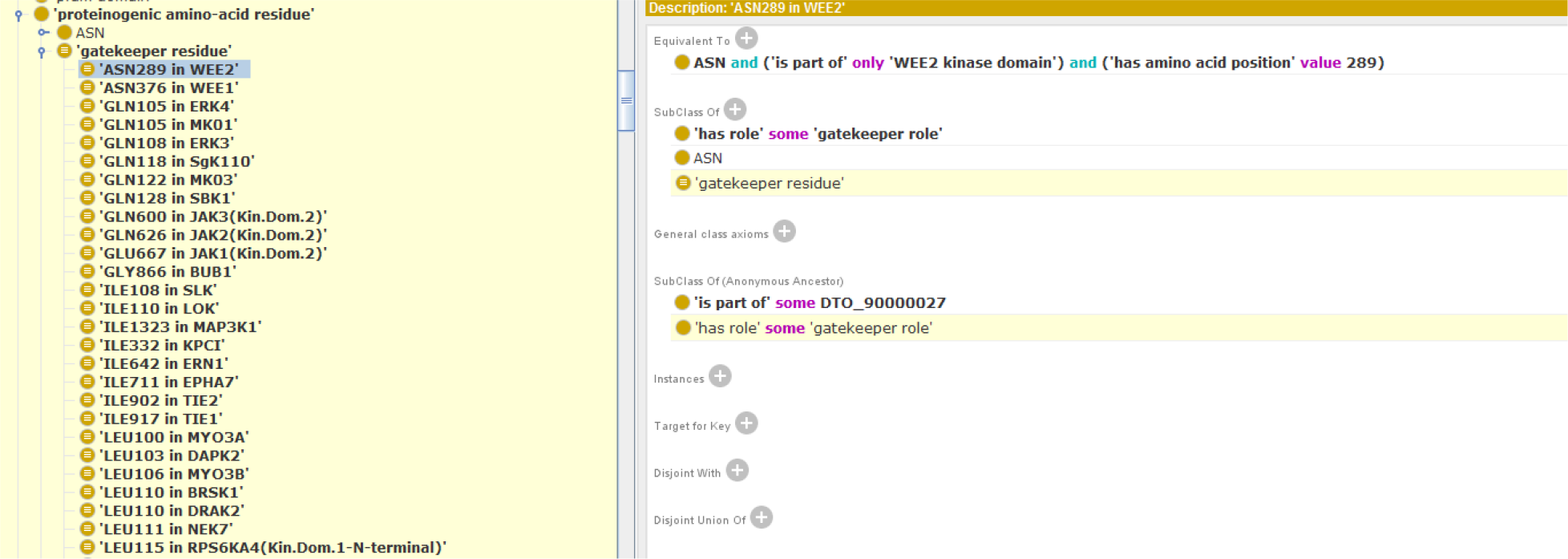
Protégé screen shot shows the inferred subclasses of gatekeeper residue.

### 3.5 DTO shines light on Tdark proteins

With integrated information about drug targets available in DTO, it is possible, for example to query information for Tdark kinases for which data in LINCS is available. Kinases in the LINCS KINOMEscan assay were annotated by their (kinase) domain, phosphorylation status, gatekeeper residue and mutations as explained above. To illustrate this integration, we conducted a simple SPARQL query to identify Tdark (kinase) proteins that have a gatekeeper annotation in DTO. The SPARQL query we use to search DTO are as following:

PREFIX rdfs: http://www.w3.org/2000/01/rdf-schema#

PREFIX rdf: http://www.w3.org/1999/02/22-rdf-syntax-ns#

PREFIX owl: <http://www.w3.org/2002/07/owl#>

PREFIX dto: <http://www.drugtargetontology.org/dto/>

select ?subject ?subject_label ?p_label ?tdl_label Where {

?subject rdfs:subClassOf ?s1.

?s1 owl:onProperty <http://purl.obolibrary.org/obo/RO_0000087>; owl:someValuesFrom dto:DTO_00000002.

?subject rdfs:label ?subject_label.

?subject owl:equivalentClass ?s2.

?s2 owl:intersectionOf ?list.

?list rdf:rest*/rdf:first ?l.

?l owl:onProperty dto:DTO_90000020; owl:allValuesFrom ?k.

?k rdfs:subClassOf* dto:DTO_61000000.

?k rdfs:subClassOf ?s3.

?s3 owl:onProperty dto:DTO_90000020; owl:someValuesFrom ?p.

?p rdfs:subClassOf* <http://purl.obolibrary.org/obo/PR_000000001>.

?p rdfs:label ?p_label.

?p rdfs:subClassOf ?s4.

?s4 owl:onProperty<http://www.drugtargetontology.org/dto/DTO_91000020>; owl:someValuesFrom ?TDL.

?TDL rdfs:label ?tdl_label

}

We found in total 378 (kinase) protein containing gatekeeper residue annotations. Of those 378 proteins, (Serine/threonine-protein kinase NEK10) is a Tdark protein, two (Mitogen-activated protein kinase 4 and Serine/threonine-protein kinase WNK1) are Tbio proteins, 320 are Tchem proteins, and 54 are Tclin proteins (Supplementary table 1). We then could look for the associated disease and tissue expression information in DTO. For example, the Serine/threonine-protein kinase NEK10 (Tdark) contains a gatekeeper residue, Thr301, which is associated with breast cancer by “weak evidence”, and expressed in liver, testis, trachea with “strong evidence”.

This way, DTO provides rich information to prioritize proteins for further study, linked directly to KINOMEscan results via the LINCS Data Portal.

### 3.6 Integration of DTO in software applications

#### 3.6.1 DTO Visualization

The drug target ontology consists of > 13,000 classes and > 122,000 links. Our visualization has two options: a) a static pure ontology viewer starting with the top-level concepts featured by a collapsible tree layout (mainly for browsing concepts) and b) a dynamic search and view page where a search-by-class user interface is combined with a collapsible force layout for a deeper exploration. Figure 7 shows an excerpt of an interactive visualization of the DTO. Users can search for classes, alter the visualization by showing siblings, zoom in/out, and alter the figure by moving classes within the graph for better visualization.

**Figure 7.**
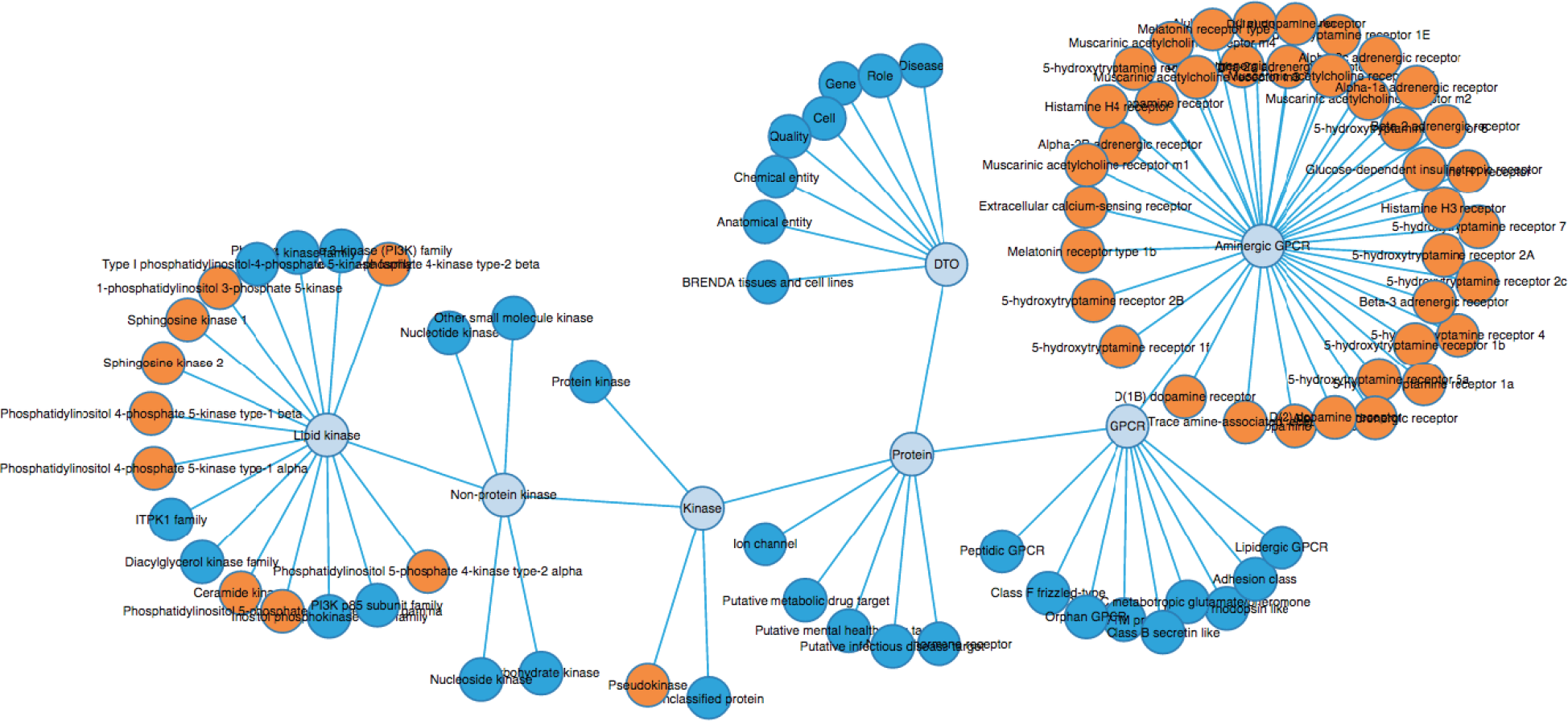
An excerpt of an interactive visualization of the DTO. The viewer is available at http://drugtargetontology.org.

#### 3.6.2 Pharos: The IDG Web Portal

Pharos is the front-end Web Portal of the IDG project (https://pharos.nih.gov). Pharos was designed and built to encourage “serendipitous browsing” of a wide range of protein drug target information curated and aggregated from a multitude of resources [11]. Via a variety of user interface elements to search, browse and visualize drug target information, Pharos can help researchers to identify and prioritize drug targets based a variety of criteria. The DTO is an integral part of Pharos; it’s user interface has been designed to integrate DTO at multiple levels of detail. At the highest level, the user can get a bird’s-eye view of the target landscape in terms of the development level through the interactive DTO circle packing visualization (https://pharos.nih.gov/dto); see Figure 8. For any suitable set of targets (e.g., as a result of searching and/or filtering), Pharos also provides an interactive sunbrust visualization of the DTO as a convenient way to help the user navigate the target hierarchy. At the most specific level, each appropriate target record is annotated with the full DTO path in the form of a breadcrumb. This not only gives the user context but also allows the user to easily navigate up and down the target hierarchy with minimal effort.

**Figure 8.**
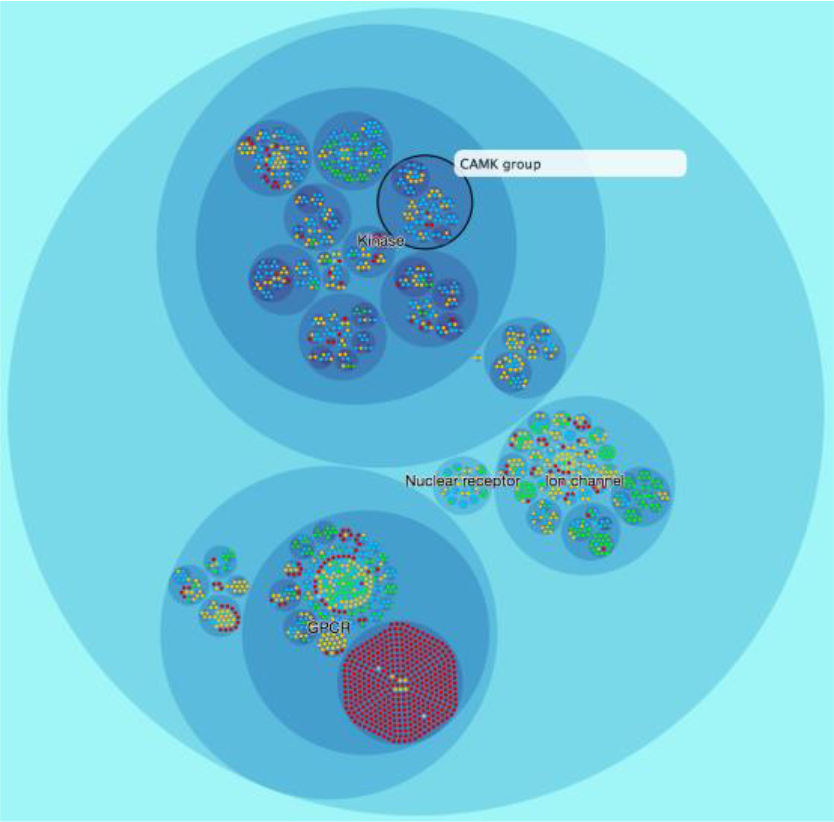
visualization of the drug target ontology using the circle packing layout available in the D3 visualization framework.

#### 3.6.3 Tin-X: Target Importance and Novelty Explorer

TIN-X is a specialized, user-friendly Web-based tool to explore the relationship between proteins and diseases (http://newdrugtargets.org/) extracted from the scientific literature [13].

TIN-X supports searching and browsing across proteins and disease based on ontological classifications. DTO is used to organize proteins and content can be explored using the DTO hierarchy.

## 4 CONCLUSION AND DISCUSSION

The IDG program is a systematic effort to prioritize understudied, yet likely druggable protein targets for the development of chemical probes and drug discovery entry points [3]. DTO covers proteins as prospective druggable targets. Druggability can be considered from a structural point of view, i.e. proteins to which small molecules can bind. This structural druggability is implicit in the selection of the IDG target families, GPCRs, kinases, ion channels and nuclear receptors for which there exist a large number of small molecule binders. Another aspect of druggability is the ability to induce a therapeutic benefit by modulating the biological function of the protein that the drug binds to. Establishing and prioritizing this functional druggability is one the main goals of the IDG project. DTO includes knowledge of protein disease association and the target development level for all proteins as a foundation to formally describes drug mechanisms of actions. DTO provides a framework and formal classification based on function and phylogenetics, rich annotations of (protein) drug targets along with other chemical, biological, and clinical classifications and relations to diseases and tissue expression. This may facilitate the rational and systematic development of novel small molecule drugs by integrating mechanism of action (drug targets) with disease models, mechanisms, and phenotypes. DTO is already used in the Target Central Resource Database (TCRD -http://juniper.health.unm.edu/tcrd), the IDG main portal Pharos (https://pharos.nih.gov/) and the Target Importance and Novelty eXplorer (TIN-X - http://newdrugtargets.org/) to prioritize drugs by novelty and importance. The search and visualization uses the inferred DTO model, including the inferred classes described in this report.

We have illustrated how DTO and other ontologies are used to annotate, categorize and integrate knowledge about kinases, including nuanced target information of profiling data generated in the LINCS project. By doing so, DTO facilitates contextual data integration, for example considering the kinase domain or the full protein, phosphorylation status or even information important for small molecule binding, such as gatekeeper residues and point mutations. As we develop DTO and other resources, we will facilitate the otherwise challenging integration and formal linking of biochemical and cell-based assays, phenotypes, disease models, omics data, drug targets and drug poly-pharmacology, binding sites, kinetics and many other processes, functions and qualities that are at the core of drug discovery. In the era of big data, systems-level models for diseases and drug action, and personalized medicine, it is a critical requirement to harmonize and integrate these various sources of information. DTO is designed to be easily extensible and integrative to other recourses, especially applying the modularization approach. The modularization approach makes it easy for developers to create, modify, expand and maintain the ontology, especially for the big scale ontologies. Several drug target-related resources have been developed, such as the ChEMBL Drug Target Slim [38], where GO annotations are available for drug targets in ChEMBL. Protein Ontology recently enhanced the protein annotation with pathway information and phosphorylation sites information [39]. Comprehensive FDA proved drug and target information is available in [3]. As a similar and complementary resource, Open Targets project (https://www.opentargets.org/) is developed by partnership between pharmaceutical companies and EBI. DTO aims to incorporate these newly developed resources, and support an integrative drug discovery data portal as a community resource for academics use.

## List of Abbreviations

**BFO** Basic Formal Ontology; **BTO** BRENDA Tissue Ontology; **ChEBI** Chemical Entities of Biological Interest; **CLO** Cell Line Ontology; **DOID** Disease Ontology; **DTO** Drug Target Ontology; **GPCRs** G-protein-coupled receptors; **IC** Ion Channel; **IDG** Illuminating the Druggable Genome; **IDG-KMC** IDG Knowledge Management Center; **IUPHAR** International Union of Basic and Clinical Pharmacology; **LINCS** The Library of Network-Based Cellular Signatures; **NR** Nuclear Receptor; **PRO** Protein Ontology; **RO** Relations Ontology; **TCRD** Target Central Resource Databases; **TDL** Target Development Level; **UBERON** Uber Anatomy Ontology

## Authors’ contributions

YL: ontology development and implementation; SM: content curation and annotation; HKM: ontology modularization and implementation; JPT: software and database tools to build the ontology; DV: kinase domain annotation, DTO curation; MF: kinase domain module, integration with LINCS data; AK: database schema and software tools to build ontology; DTN: implementation of DTO in Pharos, UI design and implementation; LJJ: integration of DISEASES and TISSUES into DTO; RG: Pharos UI design and DTO implementation; SLM: IDG database and synchronization to DTO; OU: development and implementation of DTL; VS: LINCS data processing and annotation; JD: DTO visualizer, web-interface and backend; NN: DTO database and software tools; CC: DTO website and processes; CM: software design; UV: ontology modeling and visualization; JJY: implementation of DTO in TIN-X, UI design; CGB: TDL development, TIN-X DTO integration; TO: DTL classification and annotation, UI design, content curation, project coordination; SCS: envisioned DTO, ontology development and implementation, content curation, database and software design, project coordination.

YL, SM and SCS wrote the manuscript; HKM, DTN, RG, UV, TO contributed to writing the manuscript; all co-authors approve the manuscript,

## Acknowledgements

This work was supported by grants U54CA189205 (Illuminating the Druggable Genome Knowledge Management Center, IDG-KMC) and U54HL127624 (BD2K LINCS Data Coordination and Integration Center, DCIC). The IDG-KMC (http://targetcentral.ws/) is a component of the Illuminating the Druggable Genome (IDG) project (https://commonfund.nih.gov/idg) awarded by the NCI. The BD2K LINC DCIC is awarded by the National Heart, Lung, and Blood Institute through funds provided by the trans-NIH Library of Integrated Network-based Cellular Signatures (LINCS) Program (http://www.lincsproject.org/) and the trans-NIH Big Da-ta to Knowledge (BD2K) initiative (https://datascience.nih.gov/bd2k). Both IDG and LINCS are NIH Common Fund projects. We also acknowledge resources from the Center for Computational Science (CCS) at the University of Miami.

